# Learning common and specific patterns from data of multiple interrelated biological scenarios with matrix factorization

**DOI:** 10.1101/272443

**Authors:** Lihua Zhang, Shihua Zhang

## Abstract

High-throughput biological technologies (e.g., ChIP-seq, RNA-seq and single-cell RNA-seq) rapidly accelerate the accumulation of genome-wide omics data in diverse interrelated biological scenarios (e.g., cells, tissues and conditions). Data dimension reduction and differential analysis are two common paradigms for exploring and analyzing such data. However, they are typically used in a separate or/and sequential manner. In this study, we propose a flexible non-negative matrix factorization framework CSMF to combine them into one paradigm to simultaneously reveal common and specific patterns from data generated under interrelated biological scenarios. We demonstrate the effectiveness of CSMF with four applications including pairwise ChIP-seq data describing the chromatin modification map on protein-DNA interactions between K562 and Huvec cell lines; pairwise RNA-seq data representing the expression profiles of two cancers (breast invasive carcinoma and uterine corpus endometrial carcinoma); RNA-seq data of three breast cancer subtypes; and single-cell sequencing data of human embryonic stem cells and differentiated cells at six time points. Extensive analysis yields novel insights into hidden combinatorial patterns embedded in these interrelated multi-modal data. Results demonstrate that CSMF is a powerful tool to uncover common and specific patterns with significant biological implications from data of interrelated biological scenarios.

## Introduction

With the rapid development of high-throughput sequencing technologies, numerous omics data have been generated in diverse biological scenarios, which provide unprecedented opportunities to investigate the underlying biological processes among them (1). For example, the encyclopedia of DNA elements (ENCODE) project makes a variety of ChIP-seq data of a wide assortment of cell types available; The Cancer Genome Atlas (TCGA) project generates multiple types of omics data for various cancers. Moreover, the throughput of single-cell RNA sequencing (scRNA-seq) has been significantly improved, providing a chance for comprehensively viewing the heterogeneity of cells. Integrative and comparative analysis of such data is becoming an urgent need (1). Mathematically, these genomic data can be regarded as data matrices, whose analysis method is based on matrix signal extraction and computing.

Classical data dimension reduction and pattern discovery tools such as principle component analysis (PCA), independent component analysis (ICA) and non-negative matrix factorization (NMF) are powerful techniques for analyzing high-dimensional data matrices. PCA is an effective tool for dimension reduction and visualization of such data. ICA seeks to separate such data into a set of statistically independent components. However, their decomposition results are restricted to orthogonal or independent vectors in new feature spaces and often lacks interpretability. Moreover, they are only designed for resolving one data matrix at a time. All of this limits their validity in joint pattern recognition and comparative analysis from the accumulated multiple datasets. Thus, the urgent needs for analyzing and integrating multiple data matrices prompt us to design new tools to extract hidden structures or patterns for many practical applications.

Compared to PCA and ICA, NMF not only performs dimension reduction, but also provides a better way to explain structured data. Early studies have also adopted joint non-negative matrix factorization (jNMF) and its network-regularized variant to conduct integrative analysis of multi-dimensional genomics data for extracting combinatorial patterns (2,3). More recently, an integrative NMF study further extends this framework to study heterogeneous confounding effects among different data sets (4). The aforementioned methods mainly focus on uncovering common patterns embedded in different types of data from the same biological condition. However, an unsolved valuable and urgent issue is how to perform integrative and comparative analysis on the same type of data from multiple biological conditions (e.g., different cancer types or subtypes, two or multiple epigenomic conditions) in the big data era.

A few advances have been made towards integrative and/or comparative analysis of datasets from multiple conditions. For example, differential principal component analysis (dPCA) is an efficient tool for analyzing multiple ChIP-seq datasets to discover differential protein-DNA interactions between two conditions (5). However, it only extracts differential patterns for data matrices with matched rows and columns. Tensor higher-order singular value decomposition method has also been adopted to perform integrative analysis of DNA microarray data from different studies (6). However, it was only designed for pairwise or multiple datasets (represented by tensors) that have the same row and column dimensions. Therefore, neither of these two methods can be applied to data with only one matched dimension in a unified framework. In general, data under diverse conditions may have different sizes of samples or features. ICA has been recently employed to reveal cancer-shared and cancer type-specific signals by first applying it to each cancer expression data separately (7). Simultaneous determination of common and specific patterns for the data matrices with the same row (or column) dimension among different biological conditions remains an outstanding challenge.

To this end, we propose an integrative and comparative framework CSMF to simultaneously extract <u>**C**</u>ommon and <u>**S**</u>pecific patterns from the data of two or multiple biological interrelated conditions via <u>**M**</u>atrix <u>**F**</u>actorization. CSMF can be suitable for analyzing RNA-seq, ChIP-seq and other types of data. Extensive analyses with four biological applications demonstrate that CSMF can help yield novel insights into hidden combinatorial patterns behind interrelated multi-modal data. Specifically, four applications include (I) the histone modification data for K562 and Huvec cell lines profiled using ChIP-seq from ENCODE, (II) the gene expression data of breast invasive carcinoma (BRCA) and uterine corpus endometrial carcinoma (UCEC) from TCGA, (III) the gene expression data of three breast cancer subtypes from Breast Cancer International Consortium (METRABRIC) datasets, and (IV) single-cell RNA-seq data of six time points about stem cell differentiation. CSMF is expected to be a powerful tool for uncovering common and specific patterns with significant biological implications across data of different interrelated biological scenarios.

## Results

### Problem formulation

In this study, we aimed to develop a computational method CSMF for learning common and specific patterns (or modules) among data from multiple biological conditions (Fig. 1). Here we take the gene expression data *X*_*1*_ and *X*_*2*_ of *n* genes from two conditions with *m*_*1*_ and *m*_*2*_ samples respectively as an example to illustrate our method. A common pattern in these two gene expression data is defined by satisfying the criterion “the profiles extracted from a set of columns of *X*_*1*_ and *X*_*2*_ respectively across a common set of rows has strong association or similar profiles among them”. In contrast, a specific pattern in one gene expression data (e.g., *X*_*1*_) is defined by satisfying the criterion “the profiles extracted from a set of columns of *X*_*1*_ across a set of rows has strong association or similar profiles, but not in any sets of columns of another data (e.g., *X*_*2*_) across the same set of rows”. As shown in Fig. 1, our model has a set of variables *W*_*c*_, *W*_*s1*_, *W*_*s2*_, *H*_*c1*_, *H*_*c2*_,*H*_*s1*_, *H*_*s2*_, where 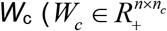, *n*_*c*_ is the common low-rank) is the common basis matrix shared by *X*_*1*_ and *X*_*2*_, *W*_*s1*_ and 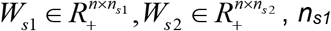, 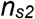 are the two specific low-ranks) are the specific basis matrices for *X*_*1*_ and *X*_*2*_, and *H*_*c1*_, *H*_*c2*_, *H*_*s1*_, *H*_*s2*_ are the corresponding common and specific coefficient matrices for *X*_*1*_ and *X*_*2*_, respectively. We then use these variables together to generate the data matrices *X*_*1*_ and *X*_*2*_. Specifically, 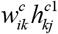 is used for the expected expression of *g_i_* in the common pattern *k*, and 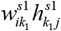 is used for the expected expression of gene *ɡ*_*i*_ in the specific pattern *k*_*1*_ for *X*_*1*_. Summing over common patterns *k* and specific patterns *k*_*1*_, the expected expression of *g*_*i*_ in condition 1 is:

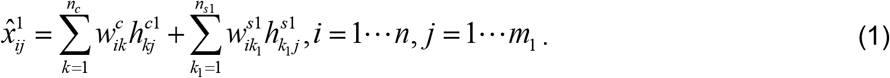

**Fig. 1.**
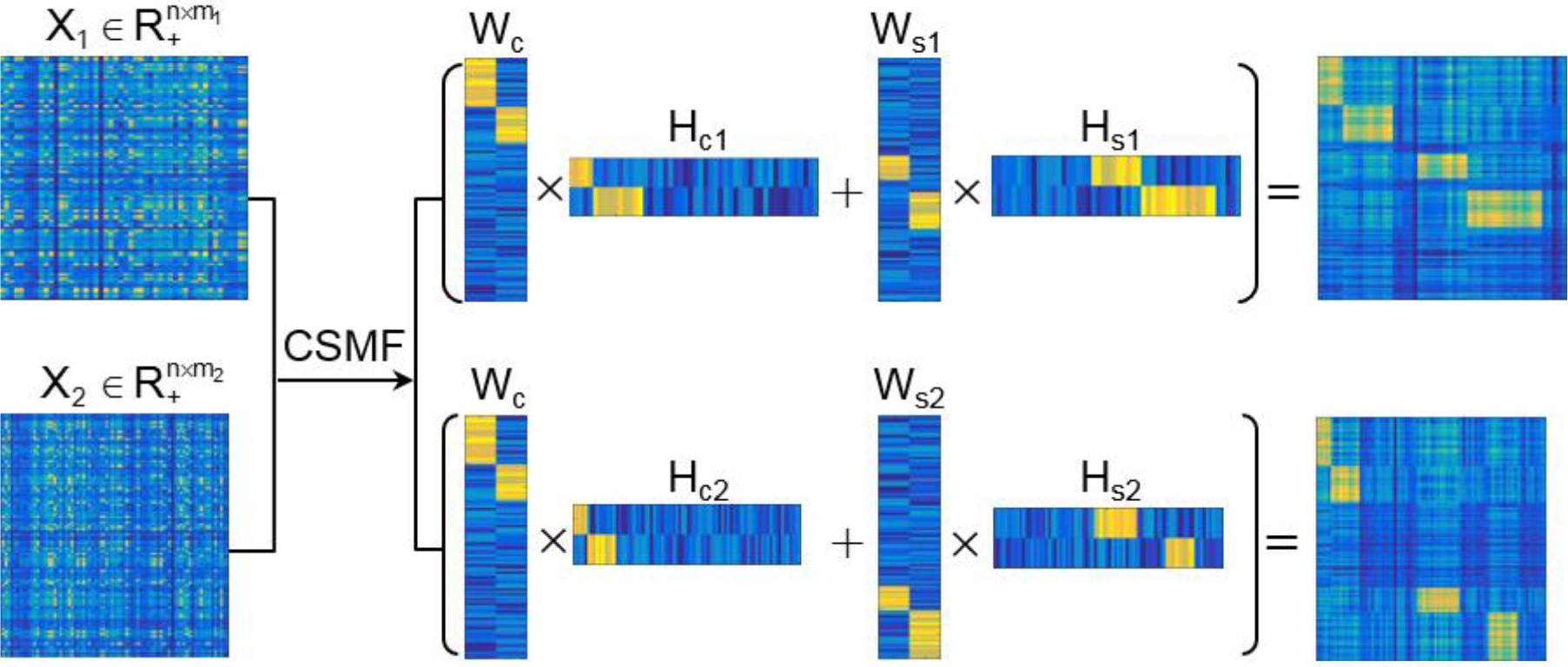
Illustration of the key idea of CSMF. *X*_*1*_ and *X*_*2*_ are the data matrices which show common characteristics with *W*_*c*_*H*_*c1*_ and *W*_*c*_*H*_*c2*_ and specific characteristics with *W*_*s1*_*H*_*s1*_ and *W*_*s2*_*H*_*s2*_. The reordered common and specific patterns in the reconstructed *X*_*1*_ and *X*_*2*_ (reordered) are demonstrated in the right panels.

Similarly, we can get the expected expression of *g*_*i*_ in the condition 2:

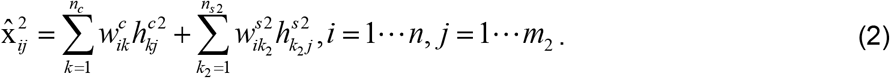

Equation (1) and (2) can be rewritten respectively in matrix format as follows:

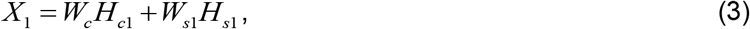

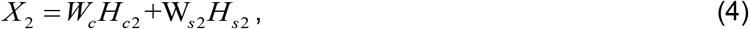

Inspired by NMF, using the squared loss function to measure the relaxation error, we can learn the variables by minimizing the following CSMF objective function:

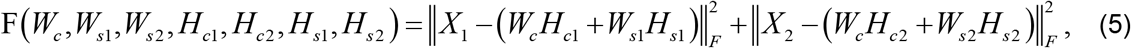

where ‖•‖_*F*_ is the Frobenius norm of a matrix, and all variables are nonnegative matrices. The two terms denote the fitting between the expected and actual expression matrices of each system. Thus, the model solution can be obtained by solving this optimization problem:

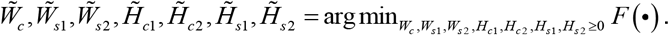

This problem can be regarded as an extension of the classical NMF, which factorizes a non-negative matrix into two low-rank ones. Naturally, the objective function *F*(•) is not convex with respect to all variables. Therefore, it is unrealistic to adopt a standard optimization algorithm to find the global minimum. To address this problem, we develop the following algorithm (**Algorithm 1**) by alternatively minimizing two sub-problems, which can be easily solved using the classical NMF algorithm. The algorithm converges to a local minimum efficiently. To obtain a robust solution, we adopted the disturbed solution of a heuristic method iNMF+ as the initial one of CSMF, and selected the solution with least collinearity (measured by Pearson correlation coefficients between any pair columns of *W*) from multiple repetitions (***SI Appendix***).

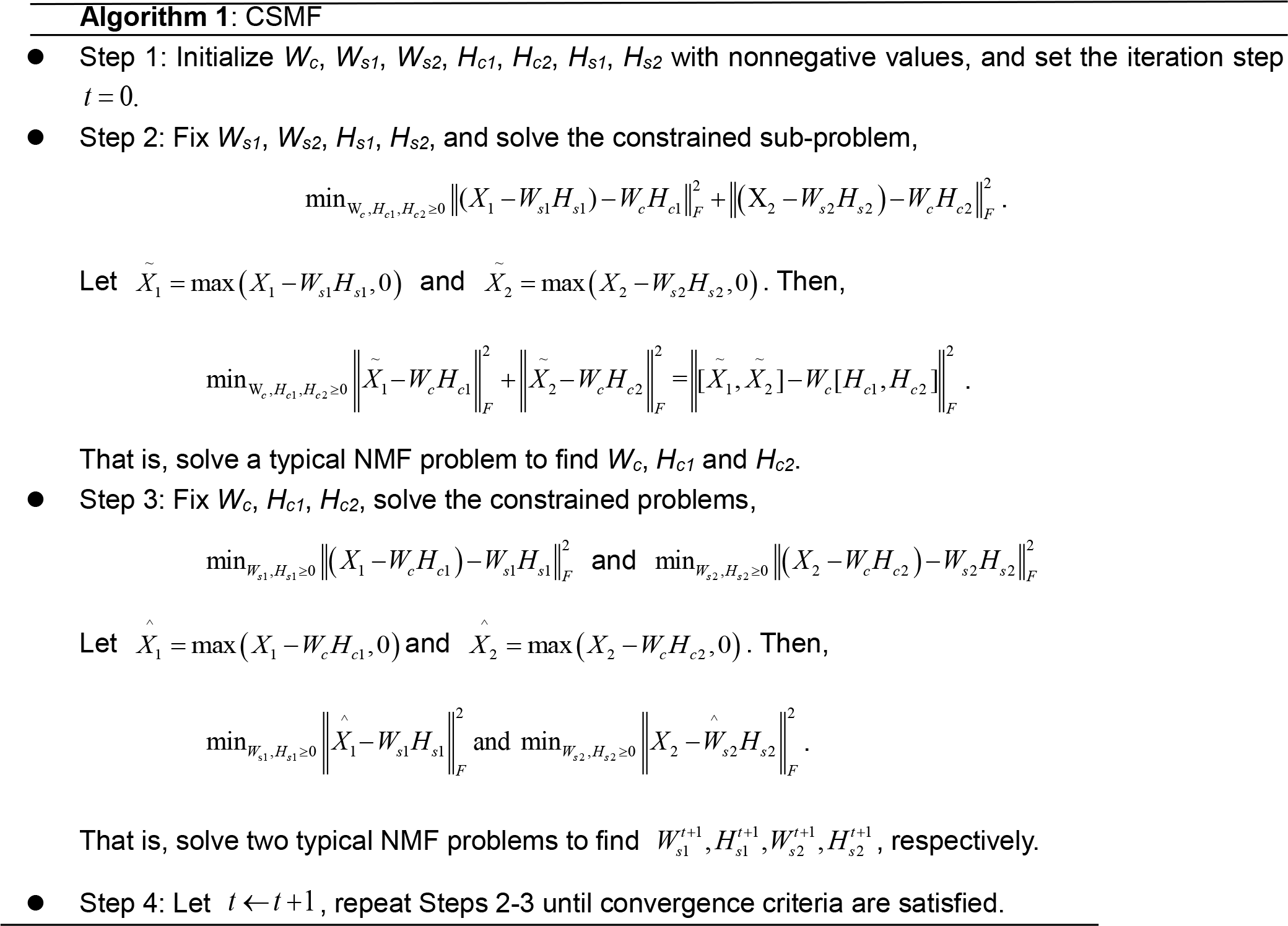

We adopt a Nesterov’s optimal gradient method to solve the typical NMF problem (NeNMF), which alternatively optimize one factor matrix with another fixed (8). NeNMF can solve the slow convergence and nonconvergence problems of other algorithms (***SI Appendix***). It is easy to see that the time complexity of CSMF algorithm is O(*T*_*o*_*T*_*i*_*nr*^2^), where *T*_*o*_ and *T*_*i*_ are the number of outer and inner iterations wherein outer iteration indicates the iteration step *t* and the inner iteration represents the iteration needed for solving any sub-problem in **Algorithm 1**, respectively, *r* is the sum of common and specific ranks, and *n* is the dimension of rows. The general form of CSMF for any number of data matrices is described in Materials and Methods. We implemented CSMF in a MATLAB package, which is available at http://page.amss.ac.cn/shihua.zhang/software.html. We first demonstrated the effectiveness of CSMF using simulation data and compared it with two naïve methods for the same purpose (***SI Appendix***).

### Determination of patterns

The obtained *W*_*c*_, *W*_*si*_ and *H*_*ci*_, *H*_*Si*_ (*i*=1,2) are used to assign both rows (features) and columns (samples) to patterns (or say modules). The maximum coefficient can be used in each column of *H*s (or each row of *W*s) to determine pattern memberships. This method restricts the assignment for each sample or feature to one and only one pattern (9). In our application, we expect one sample or feature can be assigned to multiple or none of patterns. Therefore, we employed the column-wise (row-wise) z-score of *W*s (*H*s) to determine pattern memberships as used before (3). For example, we calculated the z-score for *w*_*ij*_, which is the element *i* in the *j*-th column of *W* by 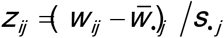, where 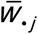 is the average value of 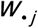 and 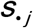 is its standard deviation. We assigned element *j* as a member of common pattern *i* if *Z*_*ij*_ was greater than a given threshold *T*.

### Application I. Determine common and specific protein-DNA interaction patterns in enhancer region between K562 and Huvec cell lines

In cellular systems, enhancers can be bound by proteins to influence gene expression by activating or repressing transcription in cells. Differential analysis of modifications of two or multiple cells are valuable to decipher their underlying distinct combinatorial and regulatory patterns. Here we demonstrate that CSMF can reveal not only differential modification patterns but also common ones. We applied CSMF to the pairwise ChIP-seq data with 58997 loci of 18 histone marks or TFs of K562 and Huvec cell lines with *n*_*c*_=1, *n*_*s1*_=3,*n*_*s2*_=3 (***SI Appendix***), and obtained *W*_*c*_, *W*_*s1*_, *W*_*s2*_, *H*_*c1*_, *H*_*c2*_, *H*_*s1*_, and *H*_*s2*_. Then we combined them to form *W* (each row represents a locus) and *H* (each column represents a mark) by

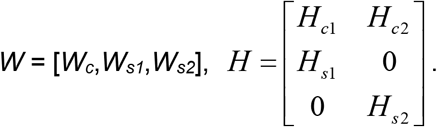

Interestingly, we can see that the common pattern (named **C**) has strong signals with CTCF mark. Coincidently, the loci of **C** pattern are significantly enriched with the motifs of CTCF (***SI Appendix***, Table S5). It is well known that CTCF is a ubiquitously expressed DNA-binding protein. Previous study suggests that CTCF-binding sites are relatively invariant across diverse cell types or cell lines including K562 and Huvec (10). Moreover, DNase mark also show strong signal in the common pattern C (Fig. 2B), which is consistent with that CTCF-binding sites co-localize with DNase I hypersensitive sites. This illustrative example demonstrates that CSMF can reveal common or shared protein-DNA interaction patterns between two cell lines, which was ignored by dPCA (5).

**Fig. 2.**
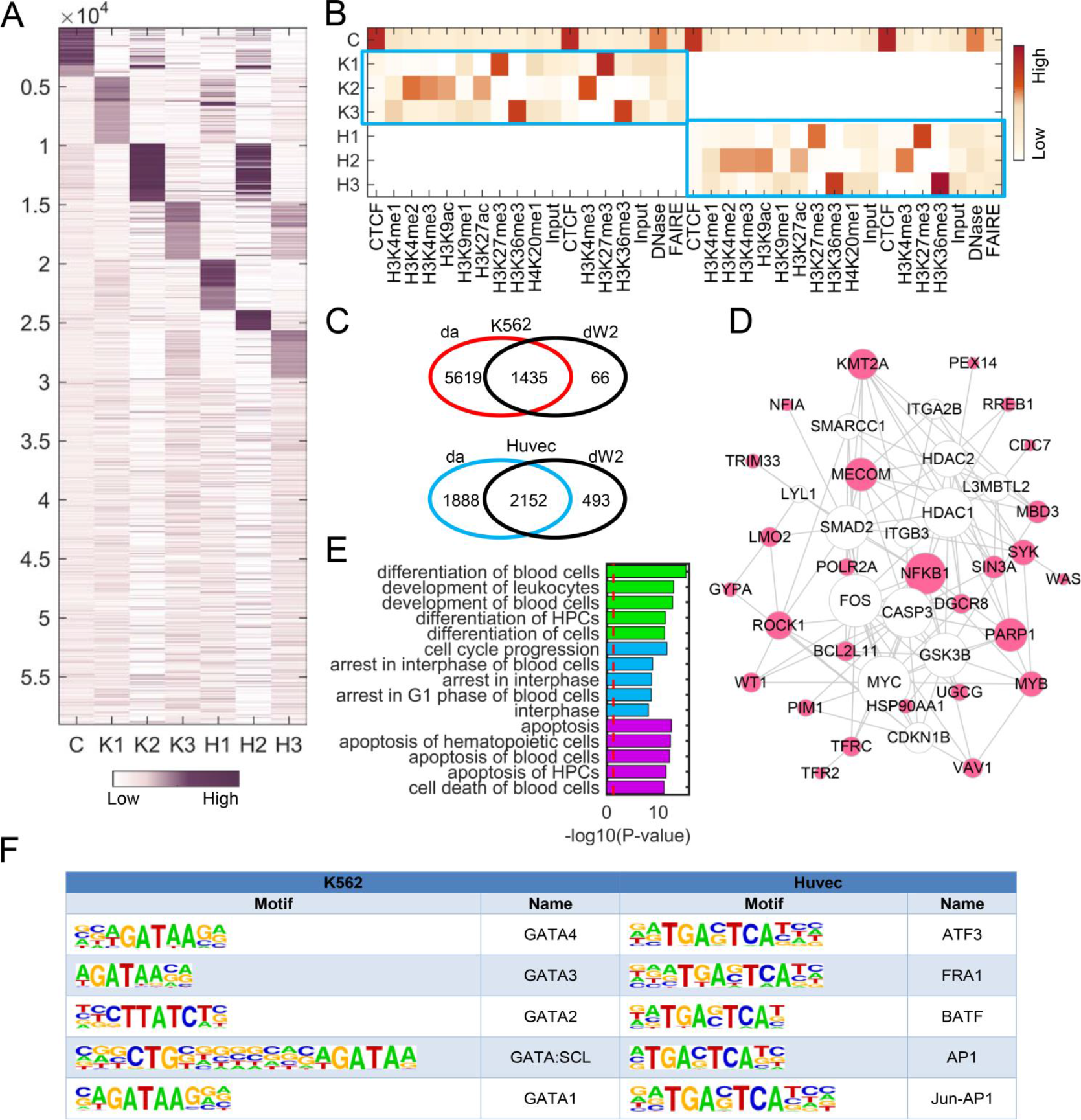
Pattern discovery by CSMF from the pairwise ChIP-seq data of K562 and Huvec cell lines with *n*_*c*_=1, *n*_*s1*_=3, *n*_*s2*_=3. (A, B) Heatmaps of the basis and coefficient matrices *W* and *H*, and *W* is reordered according to the value of its each column. Note ChIP-seq data were obtained from two different institutes (i.e., Broad Institute and University of Washington), and thus one mark may have two corresponding data. (C) Venn diagram representing the overlap loci of **da** and **dW2** across K562 and Huvec. **da** represents a locus set in which active marks have strong binding probability in K562 compared to Huvec, and vice versa detected by Wilcoxon test in the original data respectively. **dW2** represents the loci with significantly higher binding probability in K2 column of W than H2 column, and vice versa, respectively. (D) The gene network of K2 constructed using IPA and the size of gene node is proportional to its node degree. (E) The top enriched functional terms of the gene network (D) associated with cellular development (green), cell cycle (blue), and cell death (red) respectively. (F) Top 5 motifs enriched in the loci of K2 and H2 with q-value < 0.0001.

We further note that the three specific patterns (denoted as K1, K2, K3 in K562 and H1, H2, H3 in Huvec) are **marked with** diverse marks for K562 and Huvec, respectively (Figs. 2A and 2B), and the marks in K1 and K2 patterns are almost the same as marks in dPC1 and dPC2 determined by dPCA. Specifically, K1, K2, K3 are marked by strong signals of a repressive mark (H3K27me3), four active marks (H3K4me2, H3K4me3, H3K9ac and H3K27ac), and a structural mark (H3K36me3), respectively. The entries of a column in *W* represent the binding potential of marks in the corresponding pattern to the loci. Although the three patterns H1, H2, H3 are marked with the same marks as those in K562 (K1, K2, K3), the corresponding binding loci or binding intensity show distinct difference (Fig. 2A), which shapes the specificity of the two cell lines. We extracted the loci with differential binding probability (***SI Appendix***) and find that such loci in pattern K2 (**dW2**) are enriched in the differential loci with strong signals of active marks (**da**) in K562 compared to signals of active marks in Huvec detected by Wilcoxon test (Fig. 2C). Moreover, the genes closing to the loci of K1, K2 and K3 are significantly enriched in leukemia associated functions (Fig. 2E, ***SI Appendix***, Fig. S7), indicating these patterns indeed show strong specificity relating to K562. We constructed a gene functional network for each specific pattern in K562 and explore their functions by Ingenuity Pathway Analysis (IPA, http://www.ingenuity.com) (Fig. 2D and ***SI Appendix***, Fig. S7). In K1, we see a highly connected gene *RUNX1*, which is the most frequent targets of chromosomal translocations in AML, playing a critical role in leukemia development. Moreover, a previous study (11) suggested that *RUNX1* promoter are bound by EZH2 which is negatively regulated by H3K27m3, a key mark in K1. In the gene functional network of K2 (Fig. 2D), many genes (e.g., *KMT2A*, *MECOM*, *ROCK1*, *VAV1*, *BCL2L11*, *SIN3A*, *PARP1*) are related with leukemia, indicating K2 is indeed a K562-specific pattern. For example, KMT2A (also known as ALL-1 and MLL1) is a key epigenetic regulator in leukemia, which up-regulates mono-, di-and trimethylation of H3K4 (12). Moreover, many genes are connected with MYC in the gene network of K2 (Fig. 2D), which is consistent with the fact that these genes locate nearby the MYC motif. These results demonstrate that the specific binding loci of K562-specific patterns revealed by CSMF are significantly associated with leukemia.

Transcription factors (TFs) play a key role in regulating the expression of many cell line-specific genes by binding certain motifs. We note a number of significantly enriched motifs locates in the differential loci using Homer (Fig. 2F, ***SI Appendix***, Table S5). In K2, the top 5 TF motifs are all of GATA TFs, which are zinc finger DNA binding proteins, regulating transcription in cell development and cell differentiation. The functional role of most members of GATA family in leukemia has been reported in literature (13). For example, *GATA1* is a relevant biomarker for acute myeloid leukemia and its over-expression is related to the expression of *CD34* antigen and lymphoid T markers (14). *GATA2* is found in a subset of human chronic myelogenous leukemia (15) and its over-expression determines megakaryocytic differentiation (16). Overall, these results imply that GATA TFs are expected to have a higher level of expression in K562, which is consistent with K2 being a pattern of activate marks. In H2, the top 5 TF motifs are of ATF3, FRA1, BATF, AP1 and Jun-AP1 (Fig. 2F, ***SI Appendix***, Table S5), which play key roles in the development of endothelial cells (28, 29). For example, *ATF3* is highly expressed in Huvec, which protects Huvec from TNF-α induced cell death (17). *FRA1* is up-regulated in endothelial cells (18), which is consistent with that H2 is marked by active marks revealed by CSMF. Lastly, insulin/IGF pathway, PDGF signaling pathway, and VEGF signaling pathway are significantly enriched in the genes locating to the loci in H3, indicating its specificity to endothelial cells.

### Application II. Determine common and specific gene modules between BRCA and UCEC

We applied CSMF to two gene expression data of BRCA (Breast invasive carcinoma) and UCEC (Uterine Corpus Endometrial Carcinoma) and obtain five common modules (C1,C2,C3,C4,C5), two UCEC-specific modules (U1,U2) and three BRCA-specific modules (B1,B2,B3) with *n*_*c*_=5, *n*_*s1*_=2, *n*_*s2*_=3 (***SI Appendix***). Five common modules show diverse significant function enrichments with *FDR*<0.05 (Fig. 3A). These enriched biological processes relate to several key cancer hallmarks (19, 20) including cell cycle, cell division, immune response, cell death, and molecule metabolic process, suggesting their underlying common mechanisms between UCEC and BRCA.

**Fig. 3.**
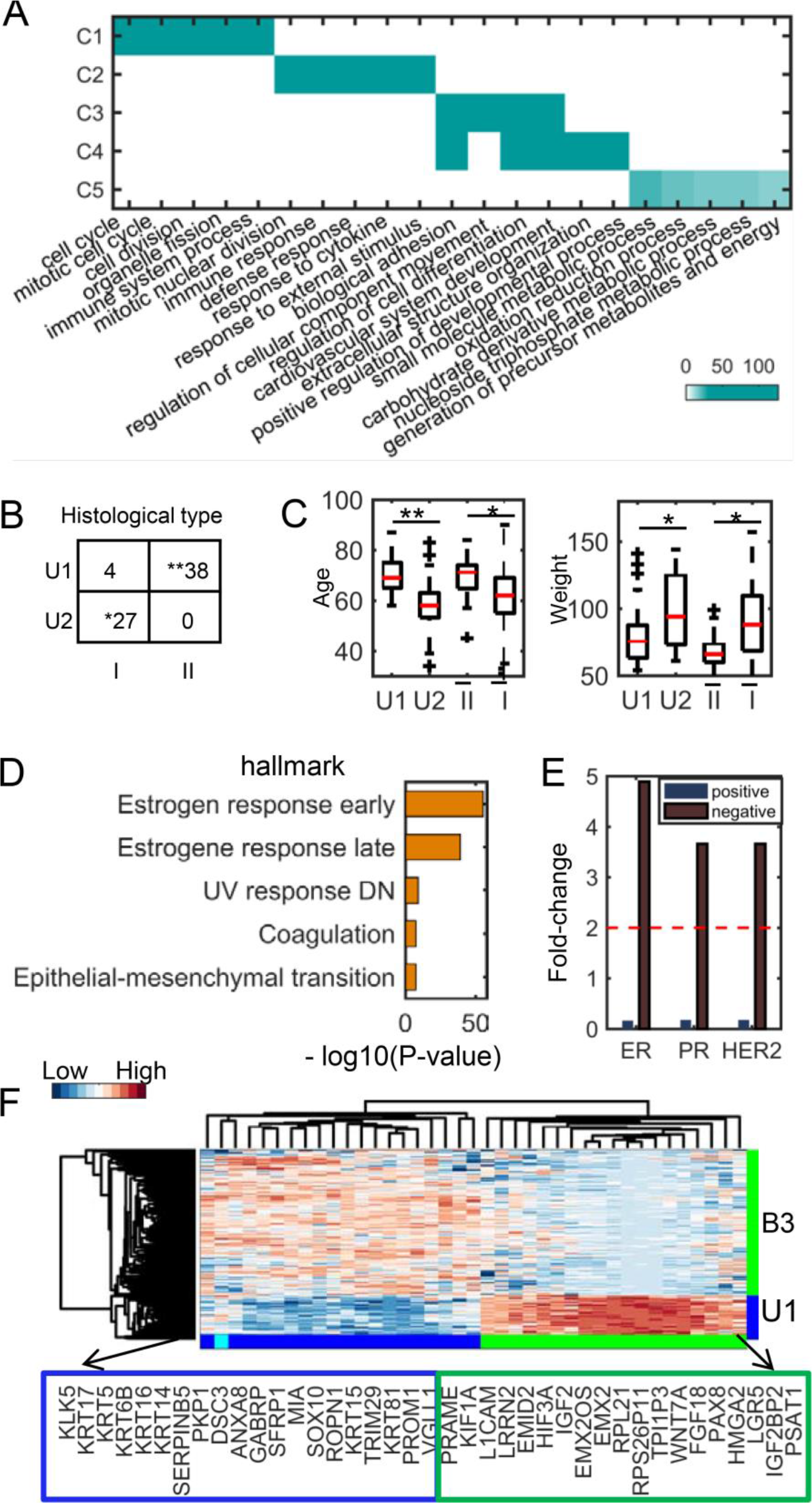
Functional and clinical analysis of UCEC and BRCA common and specific modules. (A) Functional enrichments of five common modules between UCEC and BRCA. The significance values (−log10(*q*-value)) of the enriched biological processes are shown. (B) The distribution of histological types of patients in the two UCEC-specific modules (denoted as U1 and U2). Type I and II are endometrioid endometrial adenocarcinoma and uterine serous endometrial adenocarcinoma, respectively. (C) Comparison of age and weight distribution of patients in the two UCEC-specific modules and other patients of type I and II, respectively. Ī and 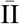 denote the type I and II patients except those in U1 and U2, respectively. (D) Top 5 enriched hallmark signatures of B1 module. (E) The Immunohistochemistry hormone receptor status enriched in B3 module. (F) The heatmap of the combined genes of top 20 highly expressed genes from U1 and B3 modules, respectively. Note *: 1e^−5^<*p*-value<0.05, **: p-value<1e^−5^.

We find that the two UCEC-specific modules U1 and U2 are significantly enriched in the two tumor histological types uterine serous carcinoma (type II) and endometrioid tumor (type I) (Fig. 3B), revealing their functional specificity as we expected. The biomarkers associated with type II carcinoma (21) such as TROP-2, kallikrein-6 and −10, claudin −3 and −4 are enriched in module U1. Intriguingly, the U1 patients are older and thinner than the U2 patients when they got sick, which is consistent with a previous conclusion about type II and type I tumors (Fig. 3C). Moreover, the age difference between patients of U1 and U2 is more significant than that between the remaining ones of type I and type II. Lastly, the estrogen receptor status of the patients in B1 are almost all positive, and top two significant hallmark gene sets enriched in this module are all associated with estrogen response (Fig. 3D, ***SI Appendix***, Table S9). In module B3, the estrogen receptor, progesterone receptor, and human epidermal growth factor receptor 2 all tend to be in negative status (Fig. 3E), implying that the module B3 is enriched with triple-negative breast tumors. Though uterine serous carcinomas share many molecular features with basal breast tumors such as high frequency of *TP53* mutation and low frequency of *TPEN* mutation, they did show some differences including distinct mutation frequency of *PIK3CA*, *PPP2R1A* and *FBXW7* (22). CSMF reveals differently expressed genes between triple-negative breast cancer in B3 module and uterine serous carcinomas in U1 module (Fig. 3F), among which, the over-expressed genes *KRT5*, *KRT6B*, and *KRT14* in B3 are indeed basal markers (23).

### Application III. Determine breast cancer subtype common and specific gene patterns

We applied CSMF to gene expression of three breast cancer subtypes including lum (denoting the combination of luminal A and B tumors), basal and her2 tumors to explore the underlying mechanisms among them with *n*_*c*_=4, *n*_*s1*_=1, *n*_*s2*_=1, *n*_*s3*_=1 (***SI Appendix***). We determined four common gene patterns (C1, C2, C3, C4) and one specific pattern (L, B, H) for each subtype using CSMF. We can see that these four common gene modules are involved in the typical cancer hallmarks like cell cycle, cell death, immune response and cellular metabolic process (***SI Appendix***, Fig. S9). More interestingly, each subtype-specific pattern tends to relate to subtype-specific pathways (Fig. 4A, ***SI Appendix***, Table S11). For example, lum-specific pattern is enriched with PID *HNF3A pathway*, where *HNF3A* (also known as *FOXA1*) is a marker of good outcome in breast cancer. Tumors with highly expressed *FOXA1* are mostly classified as luminal A ones (24). Thus, this pattern is indeed a subtype-related functional module. Basal-specific pattern B tends to be enriched in PID *Rb1* pathway and in biological processes like cell cycle, G1 phase, and G1 S transition, which is consistent with that the tumor suppressor gene *Rb1* plays a key role in regulating the cell cycle process. We note that *Rb1* deletion or mutation, *INK4a* (also known as *CDKN2A*) deletion, mutation or silencing and *CCND1*, *CDK4* and *CDK6* over-expression can cause *Rb1* loss or *Rb1* hyperphosphorylation which further disorders G1/S checkpoint (25). A previous study uncovered a strict inverse correlation between *E2F3* and *Rb1* expression in human basal-like breast cancer (27). Surprisingly, *E2F3* is highly expressed in B module, which is consistent with that *Rb1* loss is more common in triple negative breast cancers than in other subtypes (26). Interestingly, although patients with triple negative breast cancers lacking *Rb1* may have good clinical outcome under conventional chemotherapy (25), the clinical performance of patients in the basal-specific pattern B are worse among all patients in basal subtype. Therefore, the result implies that a simple loss of *Rb1* function is not responsible for the increased sensitivity of triple negative tumors to chemotherapy as suggested by previous study (26). Lastly, her2-specific pattern H tends to be enriched in *PID PLK1 pathway. PLK1* is a key regulator associated with cell cycle, and it is also associated with *her2* (28).

**Fig. 4.**
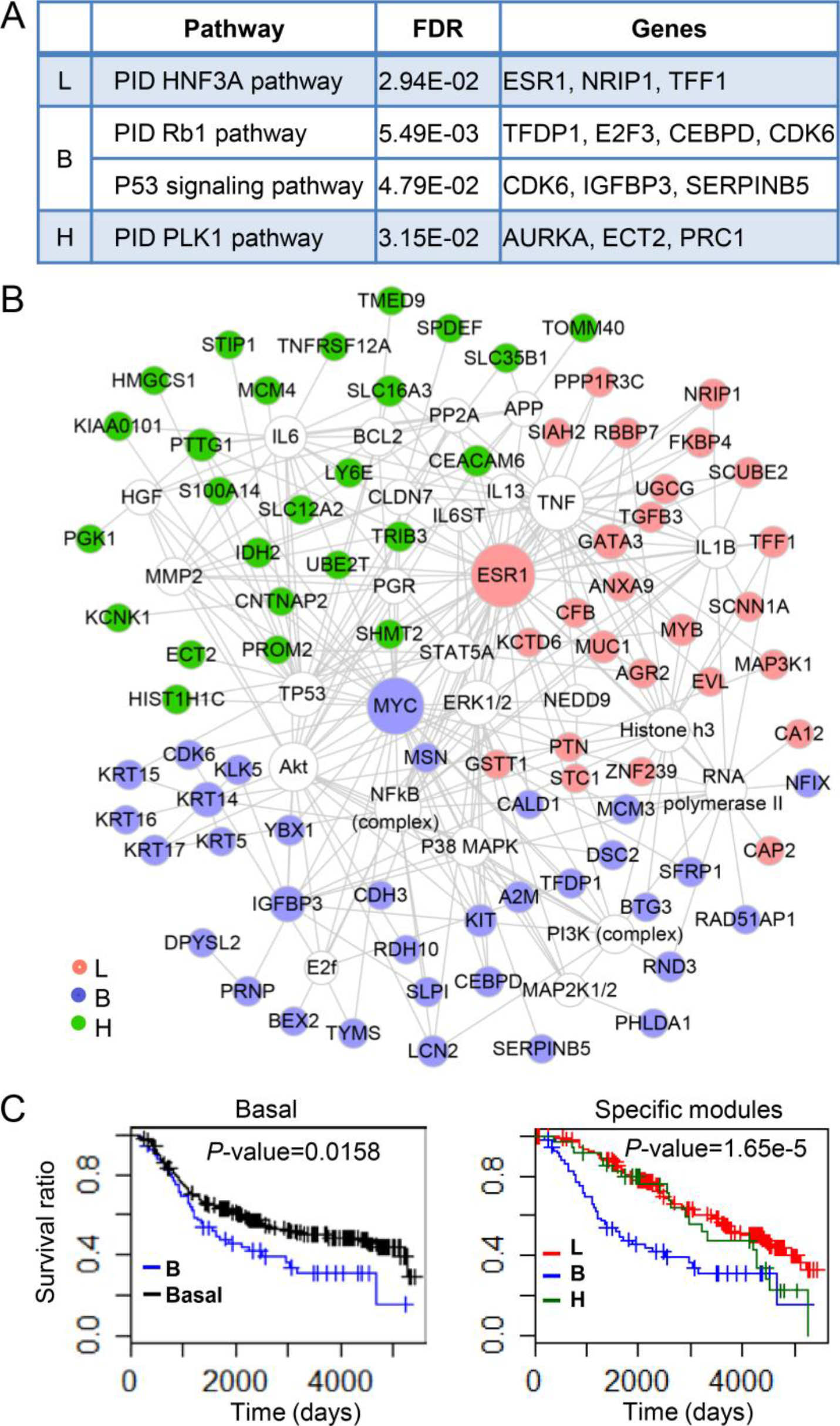
Biological functions and survival analysis of lum-, basal-and her2-specific modules (L, B, H). (A) The selected enriched pathways of each specific module. (B) Networks of genes (filled circles) in the three specific patterns visualized by Cytoscape marked with red, purple and green colors, respectively. (C) The survival curve about patients in basal-specific pattern. It is compared with that of all patients of basal subtype (left) and patients in lum- and her2-specific patterns (right), respectively.

We further constructed a gene functional network considering only the experimental verified relationships with IPA (Fig. 4B) to demonstrate the distinct functional specificity of the three subtype-specific patterns. Literature study suggests that lots of gene nodes in the network are associated with each breast subtype (***SI Appendix***, Table S12). For example, *GATA3* has been implicated in the luminal types of breast cancer, which is over-expressed in the lum-specific subnetwork. Moreover, its coding protein is a transcription factor which regulates the differentiation of luminal cells in the mammary glands (29, 30). The basal-like tumors associated basal cytokeratins (*KRT5*, *KRT6*, *KRT14*, *KRT15*, *KRT16*, *KRT17*) are highly expressed in basal specific-pattern B, demonstrating its specificity. Particularly, *KRT5* serves as an important biomarker distinguishing basal subtype and other subtypes of breast cancer. Moreover, the patients in basal-specific pattern has worse survival performance within the first five years relative to the remaining basal tumors and those in other patterns (Fig. 4C). *PGK1* is a downstream effector of her2 signaling, which contributes to the tumor aggressiveness of breast cancer. It is highly expressed in her2-specific module, confirming the functional specificity of this pattern (31).

### Application IV. Identify common and stage-specific subpopulations along the differentiation of human pluripotent cells

With the development of single cell sequencing technology, it makes possible for us to study the underlying cellular heterogeneity during human embryonic differentiation (32). We applied CSMF to scRNA-seq time-course data of six matrices consisting 8968 genes (rows) and 92, 102, 66, 172, 138, 188 cells (columns) at six time points (0h, 12h, 24h, 36h, 72h, 96h), respectively. We identified one common pattern (a cell subpopulation and its highly expressed genes) and 1, 2, 1, 1, 1, 2 stage-specific patterns at each time point (***SI Appendix***). The highly expressed genes of the common subpopulation is enriched in the biological processes including cell cycle, biosynthetic process, regulation of catabolic process, RNA processing and so on, suggesting that a cell population occurs across all time points with common functions associated with embryonic differentiation (***SI Appendix***, Table S14). On the other hand, the expression of marker genes (*POU5F1*, *T*, *CXCR4* and *SOX17*) of embryonic differentiation have nearly the same value in each subpopulation (Fig. 5A), implying each stage-specific cell subpopulation determined by CSMF is very homogenous. Moreover, heterogeneity during embryonic stem cell differentiation is always a problem for discriminating distinct phenotypic cell types. We obtained two stage-specific cell subpopulations at 12h and 96h, respectively (Fig. 5A). *POU5F1* and *T* show very diverse expression in the two subpopulations at 96h and 12h, respectively. It might suggest that one subpopulation differentiates more slowly than another one (Fig. 5A).

**Fig. 5.**
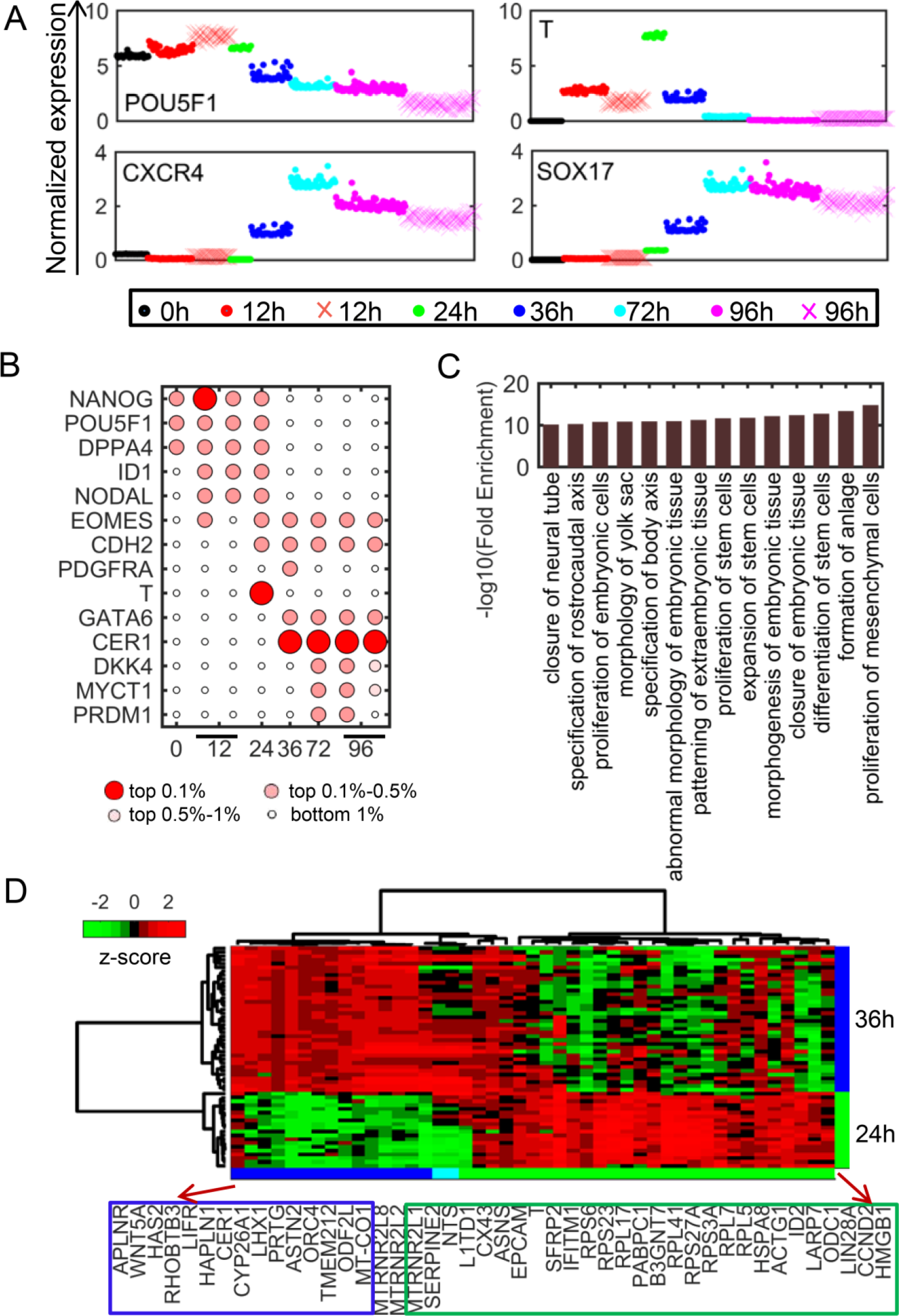
Cell subpopulations identified by CSMF from the time-course scRNA-seq data. (A) The normalized gene expression values of marker genes of cells in each specific pattern vary across time stage. Dots in specific patterns represented by different colors and dots represented by + with shallow red and rose red represent cells in 12h- and 96h-specific subpopulation II, respectively. (B) The order of significance of selected markers in each time-specific pattern (*W*). (C) The enriched function associated with embryonic development of genes, which are the intersection of genes on Wnt signaling pathway and genes differently expressed between 24h- and 36h-specific pattern. (D) The expression of genes described in (C) classify cells in 24h- and 36h-specific patterns.

CSMF can reveal subpopulation-related genes which include known differentiation-associated gene markers (Fig. 5B). For example, *NANOG* and *POU5F1* are highly expressed at 0h of differentiation. They indeed play an important role in the maintenance of pluripotency of human embryonic stem cells. While *NODAL, EOMES* and *ID1* are highly expressed at 12h of differentiation. More distinctly, *T* is highly expressed at 24h of differentiation. It is well known that *T* is the key marker of mesendoderm in embryonic stem cell studies, and it is firstly expressed in the primitive streak (33, 34). Moreover, the two definitive endoderm-specific genes *CER1* and *GATA6* are highly expressed at 36h of differentiation. Actually, *CER1* is the top important genes at 36h, 72h and 96h of differentiation (Fig. 5B). These key stage-specific gene markers revealed by CSMF are very consistent with a previous study based on a differential expression analysis tool SCPattern (32), which cannot reveal common and stage-specific cell subpopulations.

The subpopulation-related genes indeed point to key differentiation process such as the birth of definitive endoderm. To our knowledge, the hypoxic treatment experiments suggests that the birth of nascent definitive endoderm cells is a well-timed event (32). We combined the top 30 genes in 24h- and 36h-specific patterns together and obtain 45 genes after removing the ERCC family genes. This gene set is enriched in the Wnt signaling pathway, which is crucial for the development of endoderm (35, 36). These genes have different expression pattern accompanying differentiation toward definitive endoderm and are involved in differentiation of stem cells and proliferation of mesenchymal cells (Figs. 5C and 5D). Previous study suggested that cells undergo mesenchymal transition when the embryonic stem cell differentiate towards definitive endoderm during gastrulation (37). All these observations demonstrate that the pattern discovery by CSMF reveal novel hidden characteristics among the biological data of interrelated scenarios.

## CONCLUSION

With the rapid development of high-throughput technologies (e.g., ChIP-seq, RNA-seq and scRNA-seq), a huge number of genomic data of different biological conditions have been profiled and collected, providing a grand opportunity to decipher the underlying commonality and specialty among diverse biological conditions through large-scale integrative and comparative analysis. However, traditional methods fail to tackle the data reduction and comparative analysis in a simultaneous manner. To this end, we propose a powerful and flexible mathematical framework CSMF, which is suitable for analyzing data generated by different techniques such as RNA-seq, ChIP-seq and scRNA-seq. This is the first report to propose the idea to identify common and specific patterns or patterns at the same time using NMF technique.

We have demonstrated the utility of CSMF as an effective computational tool to reveal hidden patterns among complex data across diverse conditions. It is different from differential expression analysis when applied on gene expression data of case and control groups, which only identify a list of genes with statistical significance. CSMF helps us understand the biological combinatorial patterns across distinct conditions.

Selecting a well-reasoned number of common and specific patterns for CSMF is a challenging issue, and we propose a heuristic method to address it (***SI Appendix***). As the CSMF is a non-convex problem, the algorithm can only get local minimum. In future studies, we will consider to design more elaborate mathematical penalties onto the factorization to enhance the pattern discovery. Besides the four applications we conducted here, CSMF can also been applied to many other kinds of data such as copy number variation, DNA methylation and miRNA expression of different conditions, which will help us to understand the data heterogeneity and underlying patterns.

## Materials and Methods

### ChIP-seq data of K562 and Huvec cell lines

We downloaded the normalized read count data for MYC analysis example from the website of dPCA of ChIP-seq (http://www.biostat.jhsph.edu/dpca/). The MYC motif analysis dataset includes 58997 loci, 18 datasets and 70 samples in K562 and Huvec cell lines. Each locus means the extensional MYC motif site that has significant signal(s) in at least one mark or TF in either cell line. We log-transformed the binding signal with a pseudo-count 1 (i.e., log2(1+count)), and averaged the value of multiple replicates of each mark for K562 and Huvec cell lines, respectively. Finally, we got the two data consisting of 58997 loci and 18 marks in this study.

### TCGA gene expression data of UCEC and BRCA

We downloaded the level 3 gene expression data (illuminahiseq _rnaseqv2-RSEM_genes_normalized) of UCEC and BRCA on 2016-01-28 from http://gdac.broadinstitute.org/. We log-transformed the expression with a pseudo-count 1 and kept the differentially expressed genes with absolute log2 (fold change) > 2 and Benjamin-Hochberg (BH) adjusted *p*-value <0.01 between cancer and normal samples by limma (38) for UCEC and BRCA, respectively. Finally, we obtained the two gene expression data of 6621 genes across 370 UCEC and 1100 BRCA tumors, respectively.

### METRABRIC gene expression data of breast cancer

The METRABRIC dataset was accessed through Synapse (synapse.sagebase.org), which contained detailed clinical annotations such as PAM50 subtype information. We focused on the tumors of luminal A, luminal B, basal and her2 subtypes. We combined the genes which were differentially expressed between each of these subtypes and the normal-like subtype by limma with Benijamini-Hochberg (BH) adjusted *p*-value <0.01 and the absolute of log2(fold change) >0.5. We also computed the median absolute deviation (mad) value of each gene across samples in each subtype and kept the gene with mad > 0.2 in at least one subtype. We treated luminal A and luminal B as one subtype, named as lum. Finally, we got the expression of 2031 genes across 1209 lum, 328 basal and 238 her2 tumors in this study.

### scRNA-seq time-course gene expression data of human embryonic stem cells and differentiation cells

We obtained the scRNA-seq data of human embryonic stem cells and differentiation cells from NCBI’s Gene Expression Omnibus (https://www.ncbi.nlm.nih.gov/geo) with the accession number GSE75748. There are 758 total cells including 92, 102, 66, 172, 138 and 188 cells at time points 0h, 12h, 24h, 36h, 72h and 96h, respectively. The gene expression values were log-transformed with a pseudo-count of 1 and normalized by the media-by-ratio with *SCPattern* R package. We computed the variation measured by standard deviation for each gene across each time point and selected genes with z-score of the variation >1. We extracted the differently expressed genes across six time points obtained by *SCPattern* (32). In total, 8968 genes and their expression were used in this study (***SI Appendix***, Table S1).

### General CSMF

In this subsection, we introduce how to learn common and specific patterns among data from multiple biological conditions. Suppose there are *K* conditions and data *X*_*i*_ represents data matrix under the *i*-th condition. We need to determine the basis matrices *W*_*c*_ and *W*_*si*_, and coefficient matrices *H*_*ci*_ and *H*_*si*_ of each condition *i* for learning the common and specific patterns of each condition *i*. We can obtain these variables by solving the following problem:

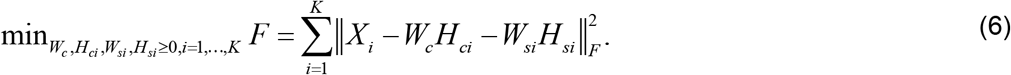

The problem (6) can be solved via a two-step procedure by solving a series of typical NMF sub-problems. In the first step, we fix *W*_*si*_, *H*_*si*_ and let *X̃*_*i*_ = max(*X*_*i*_ −*W*_*si*_*H*_*si*_,0). Then we can obtain *W*_*c*_ and *H*_*ci*_ by solving the following models,

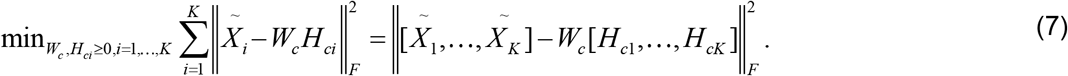

In the second step, we fix *W*_*c*_ and *H*_*ci*_ and let *X̂i* = max(*X*_*i*_ −*W*_*c*_*H*_*ci*_,0). Then we can obtain *W*_*si*_ and *H*_*si*_ by solving *K* typical NMF sub-problems and the *i*-th sub-problem is formulated as follows,

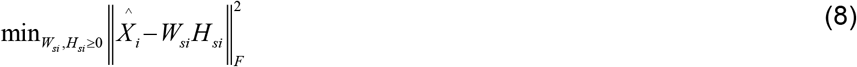

Moreover, we adopted the solution of a naive model iNMF+ as the initial one to improve the solution (see ***SI Appendix***).

### Rank selection for common and specific patterns

We chose common and specific ranks by a two-step procedure. In the first step, we took a stability-based method variant to choose rank *K*_*i*_ of each dataset and *K*_*i*_ (=*n*_*c*_+*n*_*si*_) is the sum of ranks of the common pattern and the *i*-th specific pattern. In the second step, we detected ranks of common pattern and the *i*-th specific pattern (see ***SI Appendix***).

### Tuning ranks

See ***SI Appendix***

### Simulations

We adopt a similar simulation strategy as used in (39) (see ***SI Appendix***). We applied CSMF to simulated data to demonstrate its performance and compared it with two naïve methods.

### Detailed results

More detailed analysis on the four examples were provided in ***SI Appendix***.

## Acknowledgements

This work was supported by the National Natural Science Foundation of China [No.61422309, 61621003, 61379092 and 11661141019]; the Strategic Priority Research Program of the Chinese Academy of Sciences (CAS) [XDB13040600], the Key Research Program of the Chinese Academy of Sciences, [No. KFZD-SW-219] and CAS Frontier Science Research Key Project for Top Young Scientist [No. QYZDB-SSW-SYS008].

